# A survey of naturally-occurring molecules as new endoplasmic reticulum stress activators with selective anticancer activity

**DOI:** 10.1101/2022.10.16.512420

**Authors:** Daniela Correia da Silva, Patrícia Valentão, David M. Pereira

## Abstract

The last century has witnessed the establishment of neoplastic disease as the second cause of death in the world. Nonetheless, the road towards desirable success rates of cancer treatments is still long and paved with uncertainty. With this work, we aim to select natural products that act *via* endoplasmic reticulum (ER) stress, since the latter is known to be a vulnerability of malignant cells, and display selective toxicity against cancer cell lines.

Starting from a chemical library of over 90 natural products, nontoxic molecules towards non-cancer cells were selected to be assessed for their toxicity towards cancer cells, namely the human gastric adenocarcinoma cell line AGS and the lung adenocarcinoma cell line A549. The active molecules towards at least one of these cell lines were studied in a battery of ensuing assays designed to show the involvement of ER stress in the observed cytotoxic effect, for instance, by evaluating ER stress-related gene expression or caspase activation. We show that several natural products are selectively cytotoxic against malignant cell lines, with their cytotoxicity relying on ER stress and activation of the unfolded protein response (UPR). Berberine and emodin are proposed as potential leads for the development of more potent ER stressors to be used as selective anticancer agents. Berberine was effective against the two cell models we worked with, disrupting Ca^2+^ homeostasis, inducing UPR target gene expression and ER-resident caspase-4 activation, standing out as the most promising candidate to aid in the development of novel ER stress-based strategies against neoplastic disease.

## 1. Introduction

The last few decades have witnessed an increase in favorable outcomes of cancer patients. There is, however, a long road until high success rates in cancer treatments (Miller et al., 2019). In fact, neoplastic diseases remain one of the major causes of death worldwide, second only to cardiovascular disease. Furthermore, the incidence of this type of disease has been increasing for the last few decades (Global Burden of Disease, 2015).

In the eukaryotic cell, protein synthesis and folding, as well as stress response, are inseverable of the endoplasmic reticulum (ER) function. This organelle possesses robust signaling networks that evolved towards recognition and mitigation of disturbances in proteostasis, which, globally, trigger the unfolded protein response (UPR) (Bravo et al., 2013). The UPR encompasses three major signaling branches, each activated with one of three leading sensors, namely the i) protein kinase RNA-like ER kinase (PERK), the ii) inositol-requiring protein 1 (IRE1), and the iii) activating transcription factor 6 (ATF6) (Hetz, 2012). UPR activation decreases the load of protein to be processed in the ER lumen by bringing the secretory pathway to a halt. It globally impairs protein synthesis, while enhancing ER-associated degradation of misfolded proteins (ERAD). Whenever it cannot restore proteostasis, prolonged UPR activation conducts to programmed cell death events (Bravo *et al.*, 2013; Hetz, 2012).

Cancer cells endure chronic ER stress, owing to their survival strongly relying on UPR activation since their aberrant protein synthesis rates lead to a rapid accumulation of misfolded proteins (da Silva et al., 2020). Notably, expression of UPR biomarkers is associated to multiple types of cancers, often implying therapy resistance and poor prognosis (da Silva *et al.*, 2020). The types of neoplasm in which this has been observed includes lung, breast, colon, gastric, pancreas, liver, prostate, kidney, skin, uterus and ovary carcinomas, lymphoblastic leukemias, and β-cell lymphoma (Wang and Kaufman, 2014).

Previous works have showed that when the UPR tolerance threshold is reached in cancer cells, cell death ensues. For this reason, one increasingly promising anticancer strategy is the use of molecules capable of triggering ER stress in cancer cells (Kim and Kim, 2018). Relevantly, ER stress inducers often display selective cytotoxic activity against malignant cells, which may be detrimental to the development of safer pharmacological strategies against cancer (King and Wilson, 2020). One such strategy is to induce ER stress beyond the ability of the cell to cope, leading to the occurrence of organized cell death. In fact, cell death has been shown to be elicited by all the major signaling branches of the UPR (Kim and Kim, 2018).

The natural product chemical space is prolific in ER stress and UPR modulators. Two remarkable examples are thapsigargin (Tg) and tunicamycin, commonly used as research tools to induce ER stress. Tg acts as a selective sarco/endoplasmic reticulum Ca^2+^-ATPase (SERCA) pump inhibitor, disturbing Ca^2+^ homeostasis in the cell (Thastrup et al., 1990). On the other hand, tunicamycin inhibits N-linked glycosylation of newly synthesised proteins, leading to the build-up of unfolded protein aggregates in the ER lumen (Wang et al., 2015).

Due to its immense structural variability, the natural product chemical space is regarded as a potential source of molecules that fit the criteria to be potent inducers of ER stress in cancer cells (Cheimonidi et al., 2018). Nonetheless, most reports on selective cytotoxic molecules describe cytotoxicity to non-cancer cells, though milder than towards cancer cells (Berning et al., 2019). Others are reported as selectively cytotoxic, due to their specific cytotoxicity towards few types of cancer cells, but they were not, however, compared to non-cancer cells (Kim et al., 2009).

Lung and stomach cancers represent a significant fraction of cancer incidence and death worldwide (Global Burden of Disease Cancer *et al.*, 2015). Here we report several natural products of discrete chemical classes that are more toxic against a lung cancer cell line than towards lung fibroblasts, most of them being also toxic against a gastric cancer cell line. The aim of this work was to report cytotoxic compounds against cancer cells among a library of non-toxic small molecules of natural origin that act by triggering ER stress, a known vulnerability of malignant cells (Oakes, 2017; Oakes, 2020).

## 2. Materials and Methods

### 2.1. Chemical library and reagents

The compounds 5,7,8-trihydroxyflavone, 5-deoxykaempferol, apigetrin, coumarin, cyanidin, delphinidin, diosmetin, ellagic acid, eriocitrin, eriodictyol, eriodictyol-7-*O*-glucoside, ferulic acid, fisetin, flavanone, gallic acid, gentisic acid, herniarin, homoeriodictyol, homoorientin, homovanillic acid, isorhamnetin-3-*O*-glucoside, isorhamnetin-3-*O*-rutinoside, isorhoifolin, juglone, kaempferol, kaempferol-3-*O*-rutinoside, kaempferol-7-*O*-neohesperidoside, liquiritigenin, luteolin-3’,7-di-*O*-glucoside, luteolin-4’-*O*-glucoside, luteolin-7-*O*-glucoside, malvidin, maritimein, myricetin, myricitrin, myrtillin, naringenin-7-*O*-glucoside, narirutin, oleuropein, orientin, pelargonidin, pelargonin, phloroglucinol, pyrogallol, quercetin-3-*O*-(−6-acetylglucoside), quercetin-3-*O*-glucuronide, rhoifolin, robinin, saponarin, scopolamine, sennoside B, sulfuretin, tiliroside, verbascoside and vitexin were purchased from Extrasynthese (Genay, France). The molecules (-)-norepinephrine, (+/-)-dihydrokaempferol, 3,4-dihydrobenzoic acid, 3,4-dimethoxycinnamic acid, 3-hydroxybenzoic acid, 4-hydroxybenzoic acid, berberine, betanin, boldine, catechol, cholesta-3,5-diene, cynarin, daidzein, emodin, galanthamine, genistein, guaiaverin, hesperetin, myristic acid, naringenin, naringin, *p*-coumaric acid, phloridzin, pinocembrin, quercetin-3-*O*-β-D-glucoside, quercitrin, rosmarinic acid, rutin and silibinin were acquired from Sigma-Aldrich (St. Louis, MO, USA). Caffeine was from Fluka (Buchs, Switzerland), vanillin was supplied by Vaz Pereira (Santarém, Portugal), vicenin-2 was purchased from Honeywell (Charlotte, NC, USA). Chlorogenic acid was from PhytoLab (Vestenbergsgreuth, Germany). Cinnamic acid was obtained from Biopurify (Chengdug, China). Swertiamarin was from ChemFaces (Wuhan, China).

Dulbecco’s Modified Eagle Medium (DMEM), Dulbecco’s Modified Eagle Medium/Nutrient Mixture F-12 (DMEM/F-12), fetal bovine serum, penicillin/streptomycin solution (penicillin 5000 units/mL and streptomycin 5000 μg/mL), trypsin-EDTA (0.25%), Qubit™ dsDNA HS assay kit, Qubit™ RNA IQ assay kit, Qubit™ RNA HS assay kit, SuperScript™ IV VILO™ MasterMix, and the Qubit™ Protein Assay Kit were acquired from Invitrogen (Grand Island, NE, USA). Dimethyl sulfoxide (DMSO) was acquired from Fisher Chemical (Loughborough, UK). Isopropanol was obtained from Merck (Darmstadt, Germany). 3-(4,5-Dimethylthiazol-2-yl)-2,5-diphenyltetrazolium Bromide (MTT), calcium ionophore A23187, thapsigargin, RNAzol®, chloroform, isopropanol, diethyl pyrocarbonate (DEPC), KAPA SYBR® FAST qPCR Kit Master Mix (2X) Universal, Trizma® base, sodium chloride, potassium phosphate monobasic, potassium chloride, sodium phosphate dibasic, sodium bicarbonate, D-glucose, phorbol 12-myristate 13-acetate, 4μ8C and Triton X-100 were obtained through Sigma-Aldrich (St. Louis, MO, USA). The fluorescent probe Fura-2/AM and caspase-4 assay kit (fluorometric) were purchased from Abcam (Cambridge, UK). Promega Caspase-Glo™ 3/7 Assay Kit was obtained from Promega Corporation (Madison, WI, USA). Salubrinal and JC-1 iodide were acquired from Santa Cruz Biotechnology (Dallas, TX, USA). Irestatin 9398 was acquired through Axon Medchem (Groningen, Netherlands).

### 2.2. Cell culture conditions

AGS human gastric adenocarcinoma cell line was cultured in Dulbecco’s Modified Eagle Medium (DMEM) + GlutaMAX™ containing 10% FBS and 1% penicillin/streptomycin. A549 human lung adenocarcinoma cell line was cultured in Dulbecco’s Modified Eagle Medium/F12 (DMEM) + GlutaMAX™ containing 10% FBS and 1% penicillin/streptomycin. Both were maintained at 37 °C with 5% CO_2_. In order to obtain spheroids, AGS and A549 cells were seeded in hydrophobic round-bottomed 96-well plates, in appropriate cell culture medium containing collagen (0.3 μg/mL), at a density of 5 x 10^3^ cells/well. Under these conditions, spheroids were fully formed after 72 h at 37 °C.

### 2.3. Monolayer and 3D cell viability assays

For the determination of cellular viability in monolayer cell culture, AGS cells were seeded at a density of 1.5 x 10^4^ cells/well, while A549 were seeded at 1 x 10^4^ cells/well. Afterwards, the cells were maintained at 37 °C for 24 h. After this growth period, the wells were incubated with the compounds under study (100 μL/well) for another 24 h. The medium was aspirated and replaced by fresh medium containing MTT at 0.5 mg/mL. Cells were incubated for 90 (AGS) or 120 min (A549). Finally, the MTT solution was discarded, the resulting formazan crystals dissolved in a 3:1 DMSO:isopropanol solution (200 μL/well) and the absorbance at 560 nm read in a Thermo Scientific™ Multiskan™ GO microplate reader. All experiments were performed in triplicate. Results are presented as the percentage of the control value and correspond to the mean ± SEM of, at least, three independent experiments.

To determine the viability of cells under 3D culture conditions, the compounds of interest were incubated on the spheroids for a period of 7 days. After this, the Presto Blue™ reagent was added according to the instructions from the manufacturer and incubated for 4 h at 37 °C, for both cell lines. Finally, fluorescence at 560/590 nm was read on a Cytation™ 3 (BioTek) multifunctional microplate reader.

### 2.4. Evaluation of the mitochondrial membrane potential

Cells were seeded at the previously referred densities on 96-well plates. 24 h later they were incubated with the compounds of interest for another 24 h at 37 °C. Afterwards, the medium was replaced by HBSS containing the ratiometric probe JC-1 at 7 μM. After 45 min at 37 °C, the wells were carefully washed three times with HBSS. Fluorescence was read at 485/530 nm and at 530/590 nm in a Cytation™ 3 (BioTek) multifunctional microplate reader. The effect upon mitochondrial membrane potential was inferred from the ratio F_530/590_/F_485/530_. PMA at 100 nM was resorted to as a positive control for the disruption of the mitochondrial membrane. All determinations were performed in triplicate. Results are presented as fold decrease *vs* control, representing the mean ± SEM of, at least, three independent experiments.

### 2.5. RNA sample preparation, conversion to cDNA and qPCR analysis

AGS were seeded at 1.2 x 10^5^ cells/well and A549 at 8 x 10^4^ cells/well, in 12-well plates. The plates were then kept at 37 °C for 24 h prior to the incubation with compounds of interest for other 16 h. Tg at 3 μM was used as a positive control for upregulated UPR gene expression. Cells were lysed by replacing the culture medium by 500 μL of PureZOL RNA isolation reagent in each well. The lysate was pipetted up and down several times and the PureZOL reagent transferred to the duplicate well. The process was repeated, and the lysates maintained at room temperature for 5 min.

The RNA was extracted by phase separation, by adding 100 μL of chloroform to each sample and thoroughly mixing, followed by 5 min at room temperature and 15 min centrifugation at 12,000 g, at 4 °C. The resulting aqueous phase was transferred to a new tube and mixed with 250 μL of isopropyl alcohol. Again, the mixture was left to stand for 5 min at room temperature and then centrifuged at 12,000 *g* for 10 min, at 4°C. The supernatant was discarded, and the RNA pellet washed with 500 μL of 75 % ethanol and centrifuged for 5 min at 7,500 g (4 °C). Finally, the supernatant was again discarded, and the pellet was air-dried for a few minutes and resuspended in 25 μL of DEPC-treated water.

RNA quantification was performed using a Qubit™ RNA IQ assay kit and its integrity assessed with the Qubit^TM^ RNA IQ assay kit, according to the instructions provided by the manufacturer. Conversion to cDNA was performed with the SuperScript^TM^ IV VILO^TM^ MasterMix, using 1 μg of RNA sample. The primers (**Table 1**) were designed on Primer BLAST (NCBI, Bethesda, MD, USA) and synthesized by Thermo Fisher (Waltham, MA, USA). qPCR reaction was performed with the KAPA SYBR® FAST qPCR Kit Master Mix (2X) Universal in the following thermal cycling conditions: 3 min at 95 °C, 40 cycles of 95 °C for 3 s, gene-specific temperature for 20 s (specified on **Table 1**), and 20 s at 72 °C. The reactions took place in a qTOWER3 G (Analytik Jena AG, Germany), and the results were viewed on the software supplied with the equipment (qPCRsoft 4.0). GAPDH and β-actin were selected as reference genes for expression normalization. The results correspond to, at least, four independent experiments and every reaction was performed in duplicate.

**Table 1.**
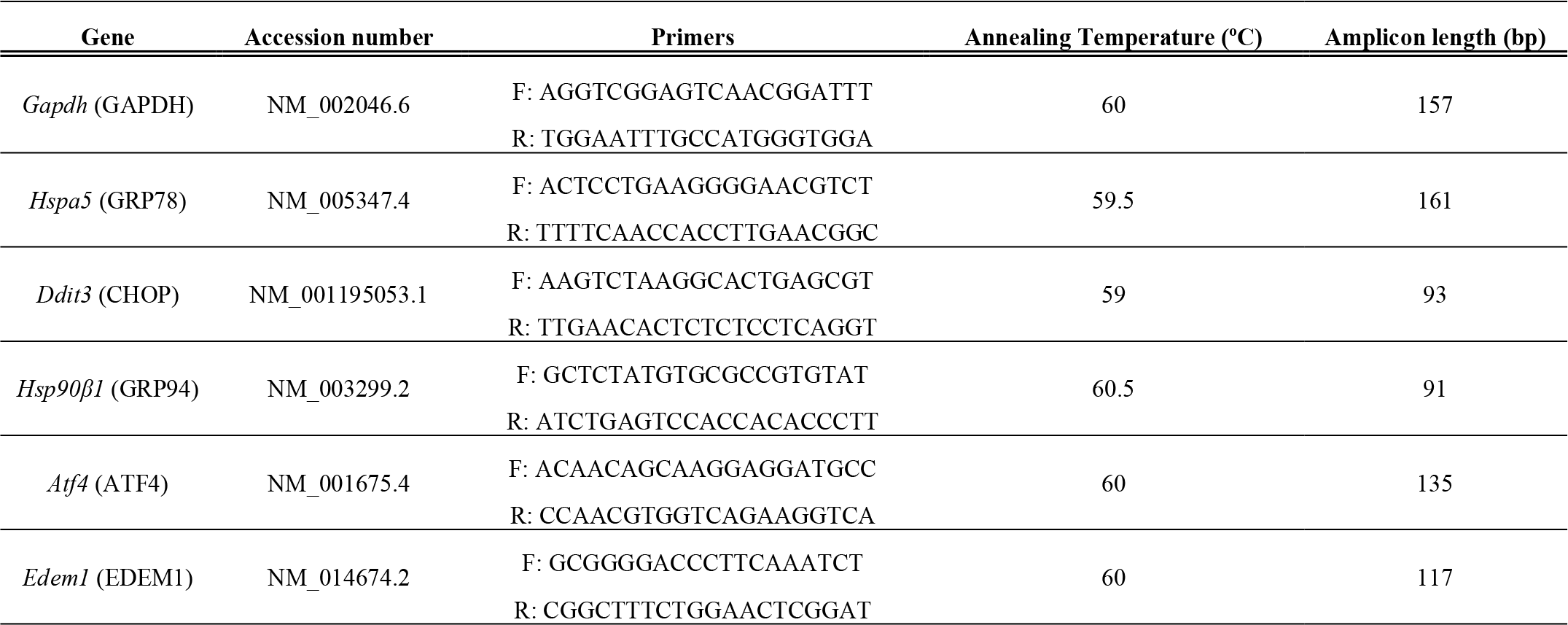
Data relative to the selected UPR-related genes and respective primers.

### 2.8. Cytosolic calcium level determination

AGS and A549 cells were plated at the aforementioned densities on black bottomed-96-well plates and incubated at 37 °C for 24 h. Then, the medium was replaced by the fluorescent probe Fura-2/AM at 5 μM in HBSS and incubated for 1 h. The molecules under study were then prepared in HBSS and added to the plate. Two hours later the wells were washed three times with HBSS and fluorescence was read at 340/505 and 380/505 on a Cytation™ 3 (BioTek) multifunctional microplate reader. Data analysis was conducted considering the ratio F_340/505/F380/505_. Each experiment was performed in triplicate. Results are presented as fold increase *vs* control and represent the mean ± SEM of, at least, three independent experiments.

### 2.9. Caspase-3 and −4 activities

To determine the activity of caspase-4, cells were plated in 6-well plates at 2.4 x 10^5^ cells/well for AGS cells and 1.6 x 10^5^ cells for A549, and incubated at 37 °C overnight. The compounds of interest were then added and cells were incubated for 6 h. After this period, cells lysates were prepared in the lysis buffer provided by the manufacturer of the caspase-4 assay kit, using a volume of 50 μL for each sample. The protein was quantified resorting to the Qubit™ Protein Assay. The assay was conducted in black-bottomed 96-well plates, using 100 μg of protein for each determination. The protein samples were incubated with the reaction buffer and the caspase-4 substrate LEVD-AFC at 50 μM, for 2 h at 37 °C and, finally, the fluorescent signal was read at 400/505 nm in a Cytation™ 3 (BioTek) multifunctional microplate reader. The experiments were individually performed in duplicate, and the results represent the mean ± SEM of, at least, three independent experiments.

The activation of caspase-3 was evaluated using the Caspase-Glo® 3/7 kit assay. Cells were seeded as described above, on white bottomed-96-well plates. 24 h elapsed, the cells were incubated with the molecules under study for a new period of 24 h. Staurosporine at 100 nM was employed as a positive control for caspase-3 activation. Afterwards, 60 of the 100 μL of medium contained in each well were discarded and replaced by 40 μL of the Caspase-Glo® 3/7 buffer, containing Caspase-Glo® 3/7 substrate. The plate was incubated for 30 min at 22 °C and the luminescent signal was measured in a Cytation™ 3 (BioTek) multifunctional microplate reader. All measurements were performed in duplicate. Results are presented as fold increase *vs* control and represent the mean ± SEM of, at least, three independent experiments.

The obtained results were normalized for the DNA amounts present after 24 h incubation with the molecules. After 24 h, cells were incubated with ultra-pure water at 37 °C, for 30 min, and subsequently frozen at −80 °C. DNA was quantified with the Qubit™ dsDNA HS Assay Kit. Caspase activity results were then normalized with the determined DNA amounts.

### 2.10. Study of the involvement of the PERK and IRE-1 branches of the UPR on cell viability

Cells were seeded as described for the MTT reduction assay and incubated with the toxic compounds in the presence or absence of salubrinal (1 μM), an inhibitor of eIF2α dephosphorylation and, thus, of PERK signaling, irestatin 9398 (5 μM), an inhibitor of the IRE-1 endonuclease activity, or 4μ8C (2 μM), an inhibitor of both endonuclease and kinase activities of IRE-1. Prior to the incubation of the molecules under study in the presence of the mentioned UPR inhibitors, a pre-incubation of 1 h with these inhibitors was conducted. The medium was then replaced by a solution containing the molecule and the inhibitor at the aforementioned concentrations. Cell viability was determined 24 h later *via* MTT reduction assay.

### 2.11. Statistical analysis

Statistical analysis was carried out on the GraphPad Prism 8 software. In order to compare single treatments with control groups, we employed the unpaired Student’s t-test, and values of *p* < 0.05 were considered statistically significant. The Grubbs’ test was used to detect outliers.

## 3. Results

### 3.1. Cytotoxicity evaluation

Considering the results obtained with 97 molecules that were devoid of toxicity towards non-cancer cells (**Table S1**, **Fig. S1A/B**), **Fig. 1 A/B** shows the molecules that caused loss of viability of the two cancer cells lines used, AGS (gastric carcinoma) and A549 (lung adenocarcinoma), at a single concentration (50 μM). The legend to the heatmap can be found in **Table S1**. Most of the hits revealed toxicity to both cells lines, with three exceptions: boldine and spermine were toxic only for AGS cells, while emodin displayed a cytotoxic effect only towards A549 cells. On the other hand, 5-deoxykaempferol, diosmetin, fisetin, berberine, genistein and kaempferol were toxic towards both studied cell lines **(Fig. 1C)**.

**Fig. 1.**
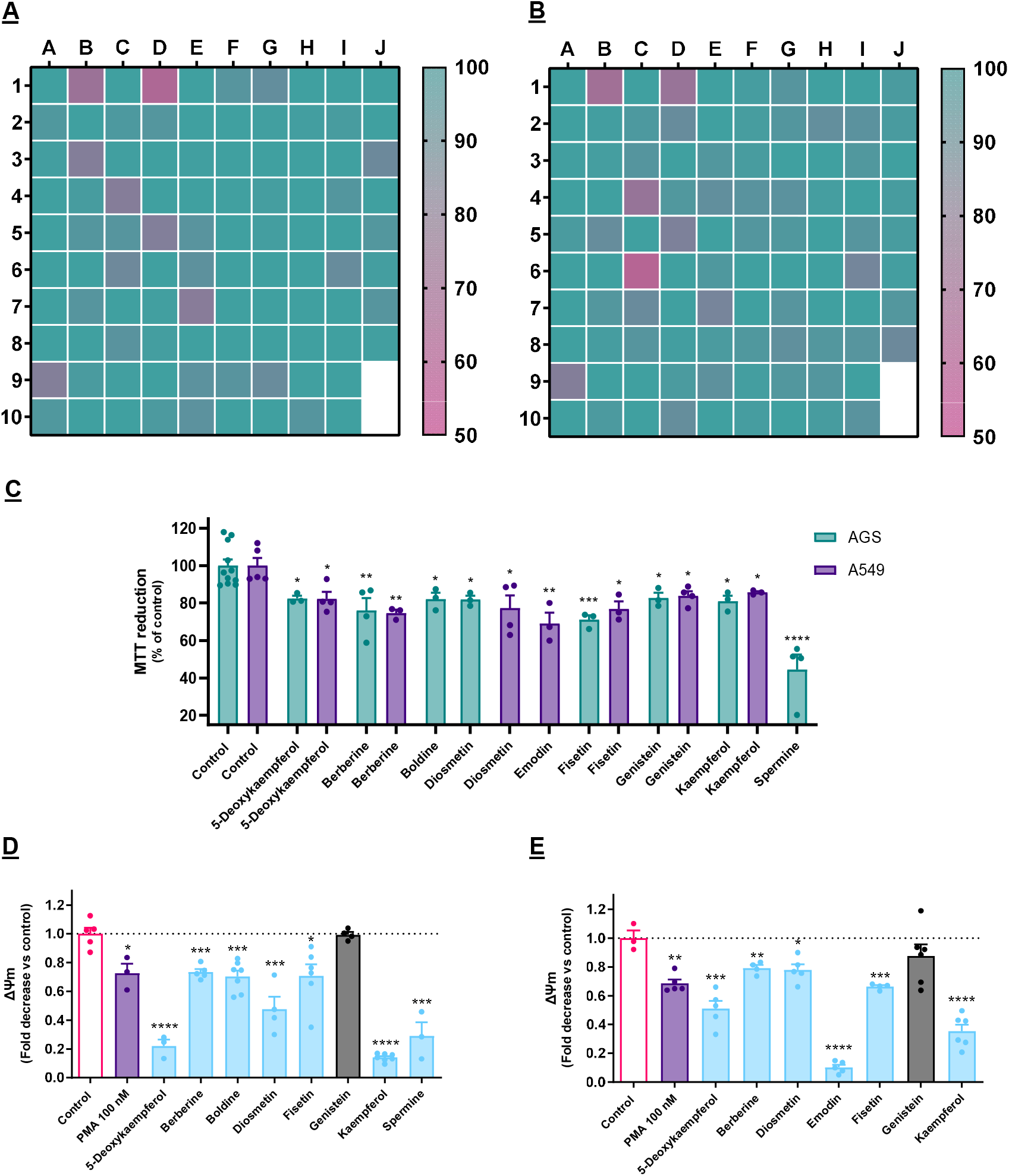
**A/B:** Impact of each compound on the viability AGS (A) or A549 (B) cells after 24 h, as determined by the MTT reduction assay. Every molecule was tested at 50 μM. Results are presented as the percentage of the control value and correspond to the mean of, at least, three independent experiments. **C:** Impact of each of the selectively cytotoxic molecules on the viability of AGS and A549 cells, representing the mean of, at least, three independent experiments ± SEM. **D/E:** Impact of the selected cytotoxic molecules on the mitochondrial membrane potential (ΔΨ_m_) of AGS (D) and A549 (E) cells. Results consider F_530/590_/F_485/530_ and express the mean ± SEM of at least three independent assays, all the latter conducted in triplicate.

As shown in **Fig. 1 D/E**, all molecules caused a reduction of ΔΨ_m_ on both cell lines, with the exception of genistein, which was for this reason dropped from the pipeline (**Fig. 1 D/E**). It is reported in the literature that PMA causes the disruption of the ΔΨ_m_ at very low concentrations, alongside increased ROS production, by being and agonist of protein kinase C (PKC), as well as inducing its mitochondrial translocation, and hence it was used here as a positive control (da Silva et al., 2017; Wang et al., 2006). We have observed that many of our active molecules at 50 μM could exert a stronger effect at this level than PMA, even though the latter was used at 100 nM. This was observed with both kaempferol and 5-deoxykaempferol on both cell lines. Furthermore, emodin displayed a potent effect on the ΔΨ_m_ dissipation in A549 cells.

### 3.2. Effect of cytotoxic molecules on UPR activation

The activation of the UPR involves the increased expression of multiple target genes, representing definitive evidence of stress upon the ER. Representative genes from multiple UPR branches and chaperones (*atf4, hspa5, ddit3, hsp90β1* and *edem1*) were selected and their expression following treatment with the cytotoxic molecules was studied.

As shown in **Fig. 2A**, both AGS and A549 cells had a similar response in terms of *atf4* and *edem1* expression when challenged with Tg, increasing the expression of these genes by around 3/4-fold. In the case of the remaining genes, namely *hspa5, ddit3*, and *hsp90β1*, the response was different between the two cell lines, A549 exhibiting a stronger response than AGS. These results show that the genes selected are adequate for assessing the UPR.

**Fig. 2.**
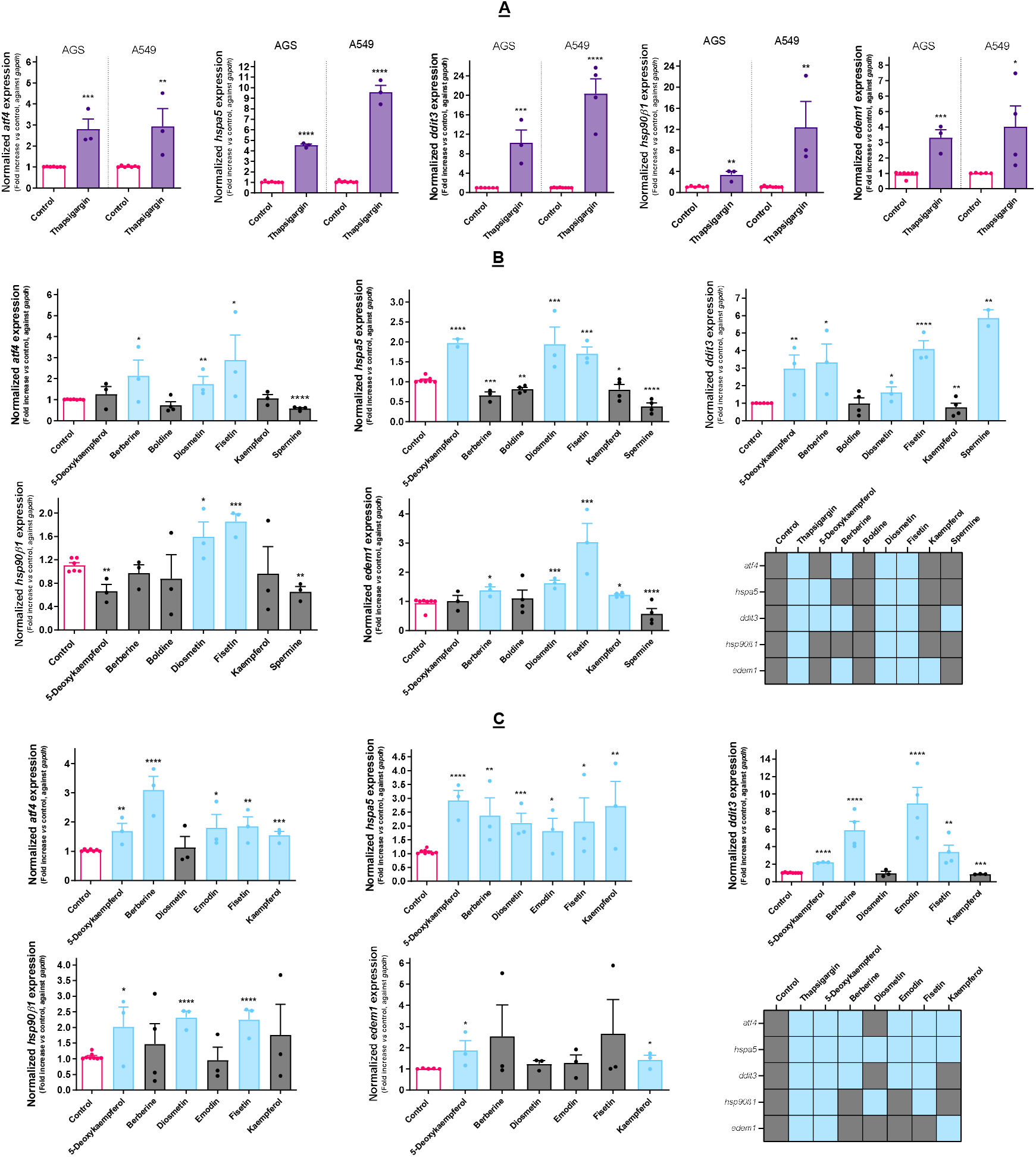
**A:** Effect of Tg (3 μM) on the expression of UPR-related genes in AGS and A549 cells, as determined by qPCR. **B:** Effect of the selected cytotoxic molecules on the expression of UPR-related genes in AGS cells, as determined by qPCR. **C:** Effect of the selected cytotoxic molecules on the expression of UPR-related genes in A549 cells, as determined by qPCR. In both cell lines, *gapdh* was selected as a reference gene for the normalization of gene expression. Results portray the mean ± SEM of three independent assays individually conducted in duplicate.

After assessing the genetic response in the context of UPR, all the molecules selected as cytotoxic were evaluated in the same conditions. As shown in **Fig. 2 B/C**, all molecules were able to upregulate the expression of at least one of the target genes, with the exception of boldine in the case of AGS cells. As so, we hypothesized that the toxicity exerted by this molecule does not involve the UPR, reason for which it was dropped from the pipeline. In these cells, diosmetin and fisetin upregulated all of the analyzed target genes, while kaempferol and spermine upregulated only *edem1* and *ddit3*, respectively. The action of 5-deoxykaempferol resulted in upregulated *ddit3* and *hspa5* expression, while berberine increased the transcription factors *atf4* and *ddit3* and also *edem1.* In what concerns to A549 cells, 5-deoxykaempferol increased the expression of all target genes, berberine and emodin increasing the expression of *atf4*, *hspa5*, and *ddit3*. Fisetin upregulated all genes but *edem1.* Kaempferol upregulated *atf4, hspa5*, and *edem1.* Diosmetin treatment resulted in increased expression of both chaperones.

### 3.3. Selected molecules impact calcium homeostasis and caspase activity

Berberine, diosmetin and kaempferol caused calcium to flow from the ER into the cytosol on the AGS cell line (**Fig. 3A**). Regarding A549 cells (**Fig. 3B**), only berberine and emodin increased the amounts of cytosolic calcium.

**Fig. 3.**
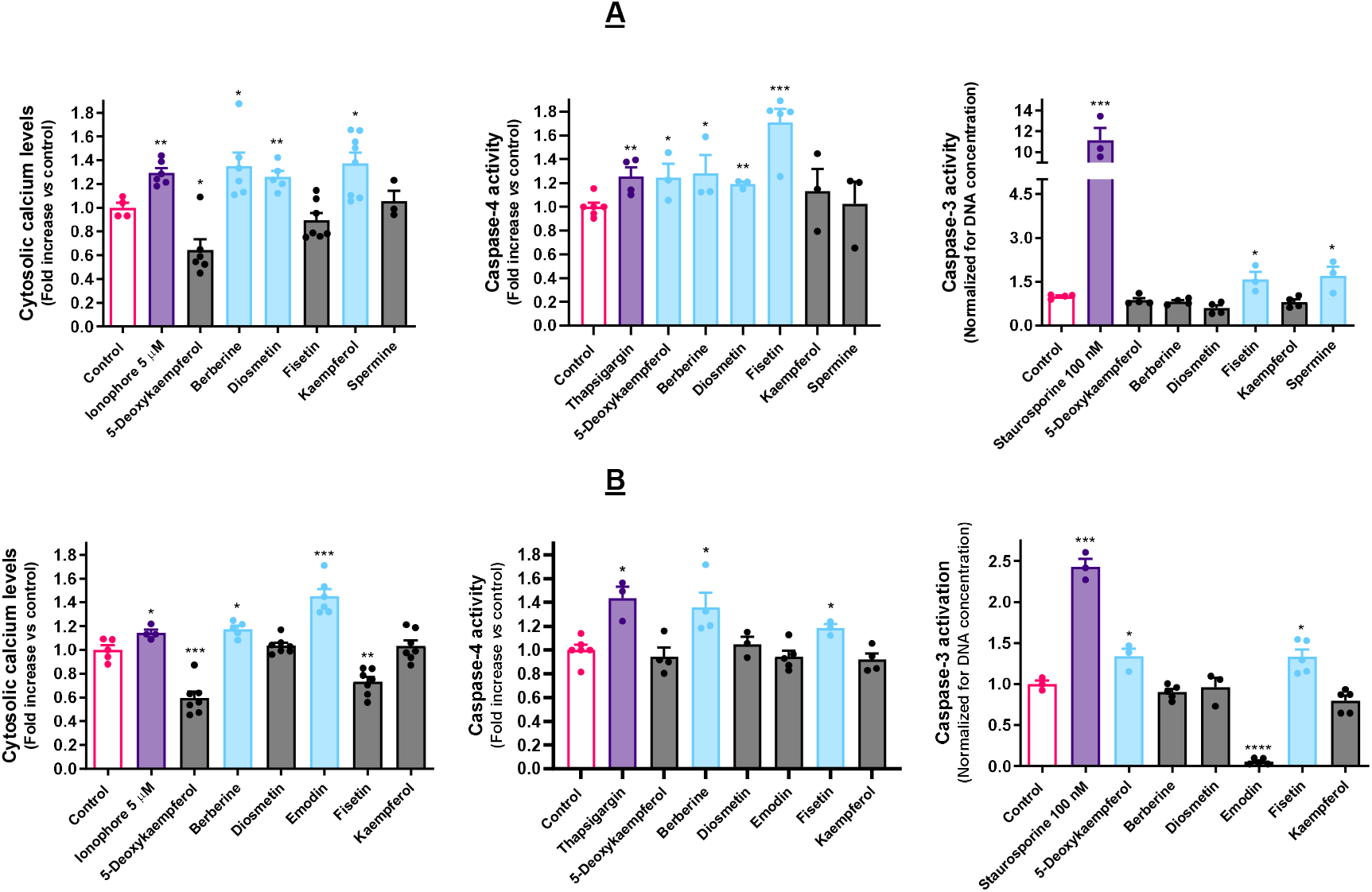
Effect of the molecules under study on the amounts of cytosolic calcium in AGS (**A**) and A549 (**B**) cells, as determined with the fluorescent probe Fura-2/AM. Results are calculated considering F_340/505_/F_380/505_ and represent the mean ± SEM of, at least, three independent assays, individually performed in triplicate. Impact of the selected cytotoxic molecules activities of caspase-4 and caspase-3 in AGS cells (**A**) and in A549 cells (**B**). The displayed results represent the mean ± SEM of, at least, three independent experiments.

Berberine elicited caspase-4 activation in both cell lines, being the only molecule that could do so in concomitance with impacting Ca^2+^ homeostasis under the selected experimental conditions. Fisetin could also significantly induce caspase-4 activation in both cell lines, while 5-deoxykaempferol and diosmetin only exerted a visible effect on AGS cells. Regarding caspase-3 activation, fisetin was the only compound that could trigger this effect in both cell lines (**Fig. 3A** and **Fig. 3B**). Other than this, 5-deoxykaempferol triggered caspase-3 activation on A549 cells (**Fig. 3B**).

### 3.4. Pharmacological modulation of UPR impacts cytotoxicity of ER stressors

In AGS cells, pre-incubation with salubrinal (1 μM) resulted in reduced toxicity of berberine (**Fig. 4A**). As for A549 cells, the same effect was observed in the cases of berberine and emodin (**Fig. 4B**). Berberine was the only of these molecules to be tested on both cells lines, and the results show that its effect extends to both. Furthermore, the salubrinal-induced increase on PERK phosphorylation enhances the toxicity of spermine in AGS cells.

**Fig. 4.**
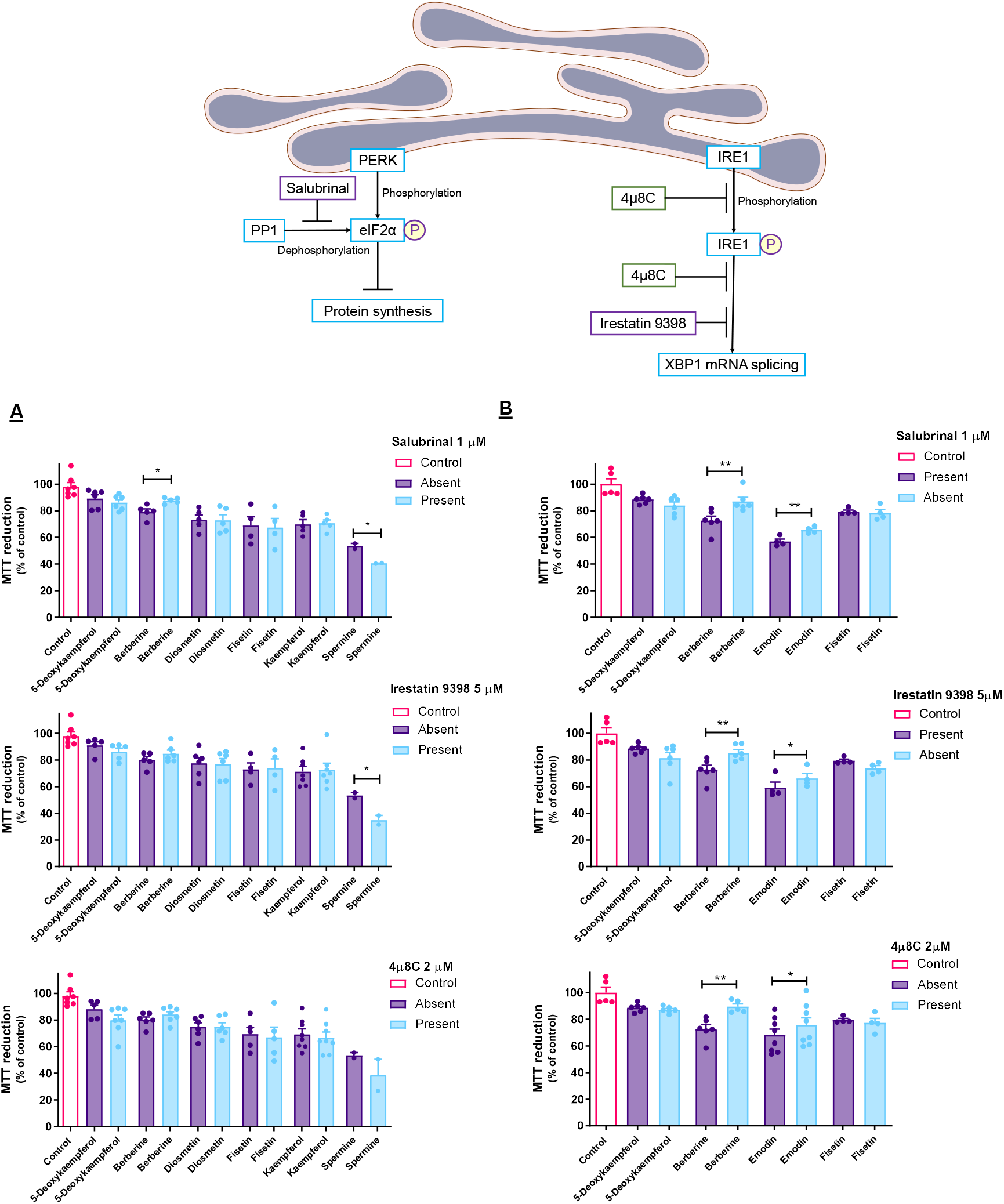
Effect of UPR inhibitors salubrinal, irestatin 9398 and 4μ8C on the impact on cell viability provoked on AGS (**A**) and A549 (**B**) cells by selected natural products. Results express the mean ± SEM of, at least, three independent experiments, all performed in triplicate.

Unlike what was verified in the presence of salubrinal, IRE1 inhibitors failed to prevent the toxicity of berberine on AGS cells (**Fig. 4A**). Regarding A549 cells, both irestastin 9398 and 4μ8C could prevent cell death induced by berberine and emodin (**Fig. 4B**).

### 3.5. Cytotoxic effect of ER stressors is retained in 3D models

A 3D cell culture system was implemented to test the selected compounds (**Fig. 5**). In AGS cells (**Fig. 5A**), berberine retained the cytotoxic effect. In the case of A549 cells (**Fig. 5B**), both berberine and emodin were active in this regard. Notably, berberine and emodin present a stronger effect than that of the positive control.

**Fig. 5.**
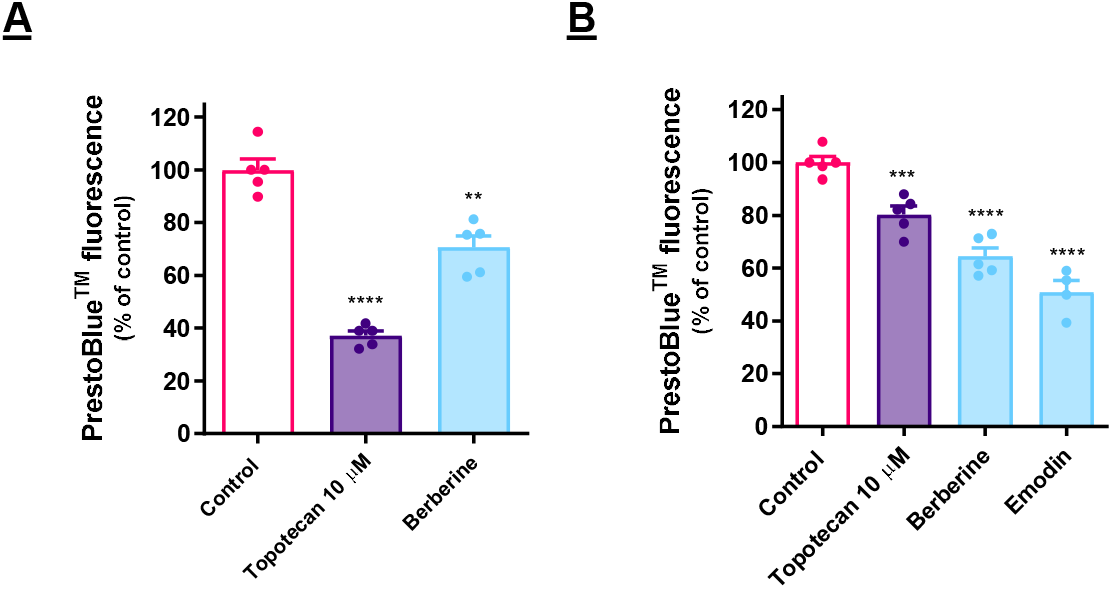
Effect of the molecules under study on the viability of AGS (**A**) and A549 (**B**) cells, in 3D cell culture conditions, as determined by the fluorescence of PrestoBlue™. Results are calculated considering F_560/590_ nm and represent the mean ± SEM of, at least, three independent assays, individually performed in triplicate.

## 4. Discussion

### 4.1. Selection of cytotoxic molecules towards cancer cells

Both the physico-chemical properties of the compound library used herein, and the information on the toxicity of the molecules in non-cancer cells, were reported in our previous work (Correia da Silva et al., 2022). As so, and in order to avoid the development of anticancer molecules lacking selectivity towards cancer cells, in the present work we have used only molecules that are non-toxic towards human non-cancer cells.

Since the molecules have shown before to be nontoxic towards a human cell line cultured from healthy tissue, we verify the selectivity of the cytotoxic effect. It is widely reported that, even though the search for molecules that do not exert undesirable side effects on healthy cells is fierce, the vast majority of attempts at discovery of such molecules has fallen short (Badisa et al., 2009).

Considering that most novel cytotoxic cancer drugs are capable of triggering an array of events that are classified as organized cell death (OCD), we were interested in further filtering the results from cell viability. We have chosen to evaluate the impact of these molecules in the mitochondrial membrane potential (ΔΨ_m_), since loss of ΔΨ_m_ is involved in most processes of OCD, its dissipation being also a hallmark of ER stress, as consequence of the upregulation of BH3-only proteins (Gupta et al., 2010).

### 4.2. The selected molecules are capable of triggering UPR activation

The *atf4* gene encodes the activating transcription factor-4 (ATF4). When the UPR is activated, PERK signaling leads to an overall decrease of the protein synthesis rate, though enhanced translation of a discrete group of proteins, that includes ATF4, occurs (Rzymski et al., 2010). In turn, ATF4 is responsible for the upregulation of genes like *ddit3*, which encodes the C/EBP homologous protein (CHOP). This transcription factor is pivotal in the process of ER stress-induced apoptosis, and its transcription may also be induced by IRE1 or ATF6 signaling (Hu et al., 2019). The *hspa5* gene corresponds to the binding immunoglobulin protein (BiP) or 78-kDa glucose-regulated protein (GRP78), while *hsp90β1* encodes the 94-kDa glucose-regulated protein (GRP94). Both of them are calcium binding proteins and the most abundantly expressed chaperones in the ER. In fact, BiP is firmly established as the major gatekeeper of UPR activation (Hetz et al., 2020). The expression of the above-mentioned genes is known to be upregulated upon ATF6 activation (Marzec et al., 2012). Under homeostatic conditions the mRNA encoding the transcription factor X-box-binding protein 1 (XBP1) is found in the cell in its unspliced form. IRE1 activation leads to its splicing and consequent upregulation of genes, such as the *edem1*, that generates the ER degradation-enhancing alpha-mannosidase-like protein 1 (EDEM1) (Kim et al., 2016). This is a well-known target of XBP1 that is associated to endoplasmic reticulum-associated degradation (ERAD) (Määttänen et al., 2010).

Tg was used as a positive control for the upregulation of the gene expression of all the aforementioned target genes in each cell line as a strategy to account for putative cell-specific differences in the cellular response, as displayed in **Fig. 2A**. This alkaloid has been extensively employed as a positive control for the occurrence ER stress and the activation of the UPR ever since its characterization as a selective SERCA pump inhibitor in the early 1990’s (Thastrup *et al.*, 1990). The SERCA pump oversees uptaking Ca^2+^ ions from the cytosol and their storage in the ER lumen, and thus its function is essential in keeping Ca^2+^ homeostasis in the cell.

The obtained results indicate that 5-deoxykaempferol, diosmetin, fisetin, berberine, emodin and kaempferol are eliciting the activation of one or more signaling branches of the UPR, reason for which they were considered for subsequent assays.

Other than this, the results make clear that UPR activation by the same molecule occurs differently in each cell model, hinting at the relevance of studying each cell type individually, since UPR activation depends on the ER capacity and function of the respective cell (van Ziel and Scheper, 2020). This is also ascertained by **Fig. 2A**, evidencing that the same Tg treatment induces different expression of target genes in each cell line.

### 4.3. ER stressors disturb calcium homeostasis and elicit caspase activation

The ER is the largest ionic calcium reservoir of the cell, being in charge of keeping appropriate concentrations of this second messenger. Tight regulation of these concentrations is essential to the upkeep of cellular homeostasis. In order to maintain it, the ER possesses an intricate network of transporters, channels, and pumps. Disturbances on calcium homeostasis, leading to its efflux into the cytosol, are a hallmark of ER stress (Krebs et al., 2015). Furthermore, the efflux of calcium ions from the ER lumen leads to the dissipation of the ΔΨ_m_ (Duchen, 2000). For all this, spatial and temporal changes in this ion are of pivotal relevance when studying ER modulators. As a positive control, we used the calcium ionophore A23187, that has been observed to trigger Ca^2+^ depletion from the ER (Li et al., 2018; Srivastava et al., 1995).

The literature shows that, even though organized cell death is inseverable from the dissipation of the ΔΨ_m_, it does not always trigger caspase activation. However, caspase activation is known as a possible downstream event of ER stress (Boya et al., 2002). Caspase-4 resides in the ER and is activated by some ER stress activators, such as Tg and tunicamycin (Hitomi et al., 2004). For this reason, Tg was used as a positive control when testing caspase-4 activation. In turn, caspase-4 can cleave procaspase-9, rendering it active and inducing the subsequent activation of caspase-3 (Momoi, 2004; Obeng and Boise, 2005). In the case of caspase-3, we used staurosporine as positive control, given that it is extensively reported to induce caspase activation (da Silva *et al.*, 2017).

The fact that molecules, such as emodin, seem to trigger every hallmark of UPR activation except for caspase activation may be due to the experimental conditions here employed in what concerns, for example, time period, since it is known that caspase expression and activation can change over time (Yang et al., 2020). Alternatively, there are reports of ER stress triggering organized cell death independently of caspase activation, which is an hypothesis for the mechanism of action of emodin in A549 cell, as well as other molecules that failed to induce caspase activation (Egger et al., 2007). Similarly, the free Ca^2+^ concentrations may be time-dependent, reason for which changes can be overlooked, and the free ions may be relocated to the mitochondria instead of remaining in the cytosol (Mekahli et al., 2011).

### 4.4. Cytotoxic molecules disturb ER homeostasis at different levels

After showing that the molecules under study were cytotoxic towards cancer cells and that this effect was mediated by the ER, we were interested in detailing the UPR branch that might have been involved, specifically the PERK and IRE1 pathways. Briefly, PERK signaling results in increased phosphorylation of the eukaryotic initiation factor 2 alpha (eIF2α), which leads global protein synthesis to a halt, while selectively enhancing the synthesis of proteins like ATF4 (Rozpędek et al., 2016). Salubrinal is a cell-permeable selective inhibitor of eIF2α dephosphorylation that acts by inhibiting protein phosphatase 1 (PP1). The effect of salubrinal protects cells against ER stress by preventing the halt in mRNA translation caused by phosphorylated eIF2α (Matsuoka and Komoike, 2015). As mentioned before, phosphorylated eIF2α selectively upregulates the expression of ATF4 that, in turn, induces the expression of target genes responsible for ER stress-induced programmed cell death mechanisms, such as *ddit3* that encodes the CHOP transcription factor (Matsuoka and Komoike, 2015; Rozpędek *et al.*, 2016). For these reasons, it is to be expected that, if the cells are undergoing events of regulated cell death related to insufficient PERK signaling, the incubation with salubrinal may relieve the cytotoxicity and allow to infer the role of this pathway in the mechanism of action of target compounds (da Silva *et al.*, 2017).

The results of berberine in co-incubation with salubrinal in AGS cells show that the toxicity of this molecule relies heavily on PERK-mediated UPR signaling, which is enhanced in the presence of this PP1 inhibitor (**Fig. 4A**). The activation of the PERK pathway may result in cell survival or programmed cell death, and, thus, in the case of the latter, it is likely that enhancing eIF2α phosphorylation resulted in increased translation of pro-apoptotic genes, and consequently, increased toxicity of spermine.

The IRE1 enzyme possesses both kinase and endonuclease activities. It is activated by transautophosphorylation and plays a dual role on the UPR signaling, being able to promote cell death or survival, like PERK. On one hand, its endonuclease activity results in the splicing of the mRNA corresponding to the transcription factor XBP1, that promotes cell survival by inducing the expression of ER-related genes. On the other hand, it can lead to cell death through increased IRE1-dependent decay of RNA (RIDD) (da Silva *et al.*, 2020; Maurel et al., 2014). 4μ8C is capable of inhibiting both activities of IRE1 by binding both its kinase and RNAse domains, simultaneously halting mRNA splicing and RIDD (Cross et al., 2012; da Silva *et al.*, 2020; Jiang et al., 2015). Irestatin 9398, on the other hand, inhibits only its RNAse domain, thus preventing XBP1 mRNA splicing (Feldman and Koong, 2007). Co-incubation of target molecules with these specific inhibitors may therefore shed a light in the involvement of IRE1 signaling on their effect.

In brief, regarding the active molecules, namely berberine towards AGS cells and berberine and emodin towards A549 cells, the selective inhibitors of PP1 and IRE1, were effective in preventing cell death. These results show that the molecules induce ER stress, leading to a collective activation of at least two of the three major branches of the UPR machinery. However, these results prove the involvement of the UPR in the effects of berberine and emodin, but they do not refute the involvement of the UPR in the cytotoxic effect of the remaining molecules, as the UPR signaling branches are often redundant and can potentially be overactivated once one of them is inhibited (Feldman et al., 2005; Ron and Walter, 2007; Vandewynckel et al., 2013).

### 4.5. Cytotoxic effect of ER stressors is retained in 3D models

Traditional monolayer cell cultures are, to this day, a powerful and the most extensively employed tool in the discovery of drug candidates and subsequent characterization of their mechanism of action. However, the surrounding environment of a cell cultured in monolayers differs considerably from an *in vivo* situation (Fennema et al., 2013). These cells attach to a surface, losing their three-dimensional conformation, lacking normal cell-to-cell interactions, and growing in the absence of extracellular matrix. The deprivation of a natural microenvironment often leads to misleading results. 3D cell culture systems recreate the natural microenvironment of the cells more closely than traditional cell cultures (Fennema *et al.*, 2013). On traditional monolayer cell cultures, the test subjects are proliferating cells, which are the ones that attach to the polystyrene surface of the cell culture plates. On the other hand, in 3D cell culture systems, the test subjects are spheroids, where the cells retain their natural morphology, cell-cell and cell-matrix interactions, and include proliferating, apoptotic and necrotic cells. The proliferating cells are present in the outer areas, whereas the core of the spheroid remains under the pressure of receiving less oxygen and nutrients, and, therefore, may be in a quiescent, hypoxic or necrotic state (Edmondson et al., 2014; Lv et al., 2017).

In the two cell models analyzed, berberine was able to induce UPR target gene activation by inducing the expression of the transcription factors ATF4 and CHOP, as well as disturbing Ca^2+^ homeostasis and promoting ER-resident caspase-4 activation. The involvement of the UPR was further confirmed upon experiments with selective pharmacological inhibitors of key UPR enzymes, and, finally, the cytotoxic effect was observed to extend to a 3D cell model. Emodin was active specifically against A549 cells, where it elicited the increase of the expression of the same genes as berberine, namely *atf4, ddit3* and *hspa5.* Moreover, it disturbed cytosolic Ca^2+^ levels and its toxicity was partly prevented by co-incubation with selective UPR inhibitors. Like berberine, it retained its cytotoxic potential in a 3D cell model.

## 5. Conclusions

The occurrence of selective cytotoxic effect among a group of over 90 naturally-occuring molecules was described and characterized. It was shown that the cytotoxicity of some of the studied molecules relies on the induction of ER stress, and we have shown their impact upon its downstream signaling pathways. In the two cell models used, berberine was able to induce UPR target gene activation, ER-resident caspase activation and disrupted Ca^2+^ homeostasis, and, for these reasons, is ascribed as the most promising compound among our chemical library to attain the goal of selective cytotoxicity against cancer cells by acting upon the ER.

## Statements and Declarations

### Author Contributions

DP has contributed to the study conception and design. Material preparation, data collection and analysis were performed by all authors. The first draft of the manuscript was written by DCS. All authors have reviewed and edited the final version of this work.

### Data availability statement

The datasets generated during and/or analysed during the current study are available from the corresponding author on reasonable request.

### Ethical approval

Not applicable.

## Acknowledgements

DCS presents special thanks to the Fundação para Ciência e Tecnologia (FCT) for the attribution of the grant (SFRH/BD/130998/2017). The work was financially supported by FCT/MCTES through national funds (project UIDB/50006/2020).

