## Supplementary Table 1 for "A survey of naturally-occurring molecules as new endoplasmic reticulum stress activators with selective anticancer activity"

**Table S1.** Legend of the heatmap, containing the molecules that were tested in cell viability assays.

| | A | B | C | D | E |
|---|---|---|---|---|---|
| **1** | Control | Berberine | Cynarin | Fisetin | Homoorientin |
| **2** | (-)-Norepinephrine | Betanin | Daidzein | Flavanone | Homovannilic acid |
| **3** | (+/-)-Dihydrokaempferol | Boldine | Delphinidin | Galanthamine | Isorhamnetin-3-*O*-glucoside |
| **4** | 3,4-Dihydrobenzoic acid | Caffeine | Diosmetin | Gallic acid | Isorhamnetin-3-*O*-rutinoside |
| **5** | 3,4-Dimethoxycinnamic acid | Catechol | Ellagic acid | Genistein | Isorhoifolin |
| **6** | 3-Hydroxybenzoic acid | Chlorogenic acid | Emodin | Gentisic acid | Juglone |
| **7** | 4-Hydroxybenzoic acid | Cholesta-3,5-diene | Eriocitrin | Guaiaverin | Kaempferol |
| **8** | 5,7,8-Trihydroxyflavone | Cinnamic acid | Eriodictyol | Herniarin | Kaempferol-3-*O*-rutinoside |
| **9** | 5-Deoxykaempferol | Coumarin | Eriodictyol-7-O-glucoside | Hesperetin | Kaempferol-7-*O*-neohesperidoside |
| **10** | Apigetrin | Cyanidin | Ferulic acid | Homoeriodictyol | Liquiritigenin |

| | F | G | H | I | J |
|---|---|---|---|---|---|
| **1** | Luteolin-3-7-*di*-glucoside | Naringenin-7-glucoside | Pinocembrin | Saponarin | Tiliroside |
| **2** | Luteolin-4'-*O*-glucoside | Naringin | Pyrogallol | Scopolamine | Trigonelline |
| **3** | Luteolin-7-*O*-glucoside | Narirutin | Quercetin-3-*O*-(-6-acetylglucoside) | Sennoside B | Vanillin |
| **4** | Malvidin | Oleuropein | Quercetin-3-*O*-glucuronide | Silibinin | Verbascoside |
| **5** | Maritimein | Orientin | Quercetin-3-ß-D-glucoside | Spermine | Vicenin-2 |
| **6** | Myricetin | *p*-Coumaric acid | Quercitrin | Sulfuretin | Vitexin |
| **7** | Myricitrin | Pelargonidin | Rhoifolin | Swertiamarin | Vitexin-2-*O*-rhamnoside |
| **8** | Myristic acid | Pelargonin | Robinin | Taxifolin | Xanthone |
| **9** | Myrtillin | Phloridzin | Rosmarinic acid | Theobromine | |
| **10** | Naringenin | Phloroglucinol | Rutin | Theophylline | |
